# Frazzled/DCC regulates gap junction formation at a Drosophila giant synapse through transcription

**DOI:** 10.1101/2025.04.08.647628

**Authors:** Juan Lopez, Jana Boerner, Kelli Robbins, Rodrigo F. O. Pena, Rodney Murphey

## Abstract

Loss of function Frazzled/DCC mutants demonstrate that the gene regulates synaptogenesis in the Giant Fiber System of Drosophila. In *frazzled* loss of function (LOF) mutants, we observe weaker physiology, characterized by longer latencies and reduced response frequencies between the GFs and the motor neurons. These physiological phenotypes are linked to a loss of gap junctions in the GFs, specifically the loss of the shaking-B(neural+16) isoform of innexin in the presynaptic terminal. A GF biophysical computational model is provided to test the role of gap junctions and the function of the Giant Fiber System. We present evidence of Frazzled’s role in gap junction regulation by utilizing the UAS-GAL4 system in Drosophila to rescue mutant phenotypes. Expression of various UAS-Frazzled constructs in a Frazzled LOF background was used to dissect the role of different parts of the Frazzled receptor in the assembly of electrical synapses. Driving Frazzled’s intracellular domain in Frazzled LOF mutants rescued axon pathfinding and synaptogenesis. This is supported by the fact that Frazzled fails to rescue the synaptic function when its transcriptional activation domain is disrupted, as shown by the deletion of the highly conserved intracellular P3 domain or by a construct with a point-mutation in the highly conserved P3 domain known to be required for transcriptional activation. The present work is the first to show how a guidance molecule regulates synaptogenesis through transcriptional regulation of synaptic components.

## INTRODUCTION

Various ligand-receptor pairs are involved in axon guidance and pathfinding; more recently, some have been shown to have a second function during synaptogenesis. Frazzled/DCC is one such class of receptor. It belongs to the immunoglobulin (Ig) superfamily of transmembrane-associated proteins defined by the Ig domain consisting of ∼100 amino acids arranged in the extracellular space (Faissner, 2009). Frazzled belongs to a specific Ig subfamily consisting of its human homologs, DCC and neogenin (Kolodziej et al., 1996). The Netrin-Frazzled pair has been shown to be involved in synapse formation in worms (Colon-Ramos et al., 2007) and flies (Orr et al., 2014).

The Giant Fiber System of Drosophila is one of the animal kingdom’s most extensively studied synaptic circuits (Allen et al., 2006; Blagburn et al., 1999; Power, 1948; Thomas & Wyman, 1983). Each Giant Fiber (GF) has a cell body and dendrites in the brain and extends a large axon out of the brain along the midline to the mesothoracic neuromere where it forms a mixed electrical-chemical synapse to the tergotrochanteral motoneurons (TTMn), resulting in a reliable, high-speed circuit that signals to the jump muscles, establishing the rapid escape behavior of flies (Allen et al., 2006; von Reyn et al., 2014). This synapse has been studied since the early 1980s, and the invertebrate gap junctions (GJs) were discovered and characterized there (Blagburn et al., 1999; Phelan et al., 2008; Thomas & Wyman, 1983). The mixed synapse is comprised of a chemical component of that releases acetylcholine (Allen & Murphey, 2007; Hernandez-Nunez et al., 2014) and an electrical component is mediated by shaking-B innexin gap junctions (Jacobs et al., 2000; Mark & King, 1983; Phelan et al., 1996, 2008; Thomas & Wyman, 1983). Shaking-B gap junctions are composed of four protein subunits in the presynaptic and four in the postsynaptic membranes (Oshima, 2020; Phelan et al., 2008). Alternative splicing of the innexin gene *shaking*-*B* produces three different innexin proteins in the Giant Fiber System: shaking-B(lethal), shaking-B(neural), and shaking-B(neural+16) (Phelan, 2005). The shaking-B(neural+16) innexin is found in the presynaptic GFs, while the shaking-B(lethal) isoform is implicated in the TTMn.

### Axon guidance molecules and synapse assembly

Netrin is a well-characterized and conserved axon guidance cue that was the first of its kind to be discovered (Boyer & Gupton, 2018; Mitchell et al., 1996; Serafini et al., 1994). The Netrin receptor Frazzled and its vertebrate homologs DCC and neogenin are known to play roles beyond axon guidance, including synaptogenesis, regeneration, and cell migration (Colon-Ramos et al., 2007; Flores, 2011; Ylivinkka et al., 2016). Previous work done in our lab suggests that Netrin-Frazzled binding regulates gap junctions in the GFs (Orr et al., 2014).

The Frazzled gene is located on the second chromosome (Kolodziej et al., 1996), and the receptor Frazzled canonically binds Netrin on growth cones via its extracellular domain. Recently, a novel, non-canonical, Netrin-independent role of Frazzled was found, where the intracellular domain (ICD) of Frazzled was shown to function as a transcriptional activator of the gene *commissureless* (in the eagle neurons) of embryonic *Drosophila* (Neuhaus-Follini & Bashaw, 2015; Zang et al., 2022). Further exploration of Frazzled’s ICD revealed that the P3 domain was responsible for this transcriptional effect. We hypothesize that a similar effect occurs in the GFs, where Frazzled may be regulating innexin GJs in the GFs at the transcriptional level.

While little is known about Frazzled function beyond canonical axon guidance and the transcription of *commissureless*, here we present evidence for the transcriptional role of Frazzled in synapse formation, structure and function. Using UAS-GAL4-driven Frazzled constructs, we dissect the molecular requirements of Frazzled for gap junction transcription in the CNS of adult *Drosophila*. We demonstrate that Frazzled’s ICD is necessary to produce the GF-specific gap junctions shaking-B(neural+16) by quantifying changes in physiology and gap junction production across our various rescue experiments.

## METHODS

### Fly stocks

*Drosophila melanogaster* were raised at 25°C, and data were collected from both male and female flies. Adults 2-4 days old were selected for testing. We generated a frazzled loss of function (LOF) mutant using *fra^3^* and *fra*^4^ null alleles (Kolodziej et al., 1996; Zang et al., 2022). When either allele is homozygous, the mutant is lethal; however, the trans-heterozygous mutant (*fra*^3^/*fra*^4^) has a low survival rate, and we used these escapers for experiments. The UAS-GAL4 system was used to fluorescently label the Giant Fibers (GFs) using the R91H05::GFP-GAL4 as our GF-GAL4 driver (Figure 1A, 1C; Jenett et al., 2012). The tested fly lines are listed in Table 1. The following parent lines were also generated and tested (Data not reported): w; *fra^3^/*CyO; R91H05::GFP-GAL4/+, w; *fra^3^/*CyO; R91H05::GFP-GAL4/Tb, w; *fra^4^*/CyO; R91H05::GFP-GAL4/Tb, and w; *fra^4^*/Gla, Bc; R91H05::GFP-GAL4/TM3. The following published transgenic lines were used in this study: UAS-FraICD-myc, *fra*^3^/CyObb; UAS-Frazzled-Myc/TM2, UAS-HA-FraE1354A, UAS-HA-FraΔP1, UAS-HA-FraΔP2, and UAS-HA-FraΔP3 (Figure 1I; Garbe & Bashaw, 2004; Neuhaus-Follini & Bashaw, 2015).

**Figure 1.**
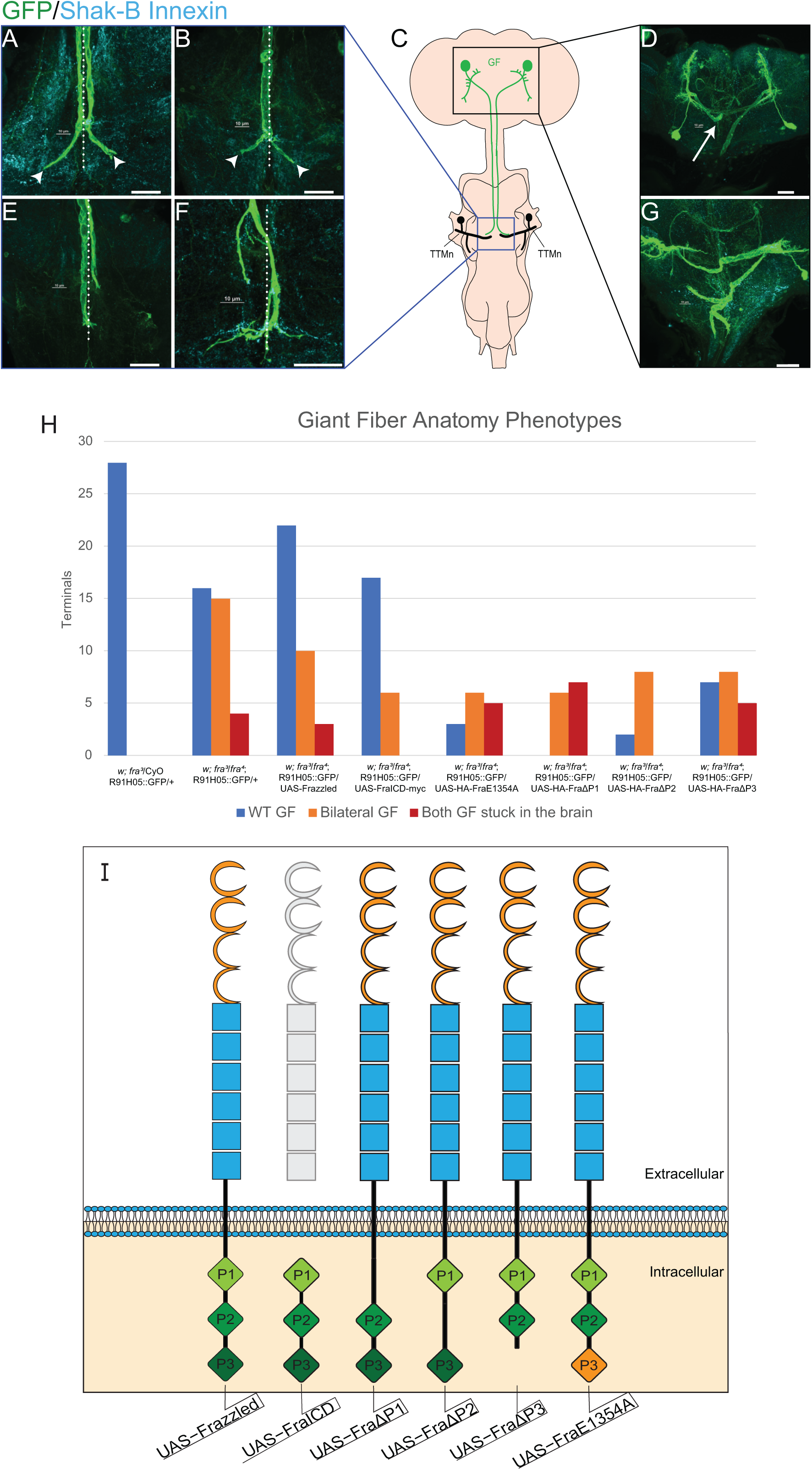
Axon guidance phenotypes and resulting physiology. A-G: Confocal images of the various axon guidance phenotypes found in frazzled LOF specimens. Midlines are shown in dotted lines. **A)** Wild-type appearing Giant Fibers and terminals (arrowheads). **B)** A single right GF forms a bilateral terminal (arrowheads). **C)** Schematic of the GF system. The GF somata are labeled green, and the axons grow along the midline to form terminals that synapse onto ipsilateral TTMn partners in black. **D)** The missing left GF from B can be seen in the brain (arrow), but its axon does not leave the brain. The right GF extends an axon that is seen in B. **E)** Rarely occurring GFs that do not form traditional bends towards their TTMn partners. **F)** Terminals in GFs driving UAS-HA-FraΔP2. **G)** neither GF grows out of the brain. **H)** The frequency of different axon guidance phenotypes for each genotype tested. **I)** Diagram showing the structural differences between the UAS constructs used in our experiments. The extracellular domain for each construct contains four immunoglobulin C2 repeats (orange) and six fibronectin III repeats (blue), while the intracellular domain consists of the P1, P2, and P3 domains (green). UAS-FraE1354A contains an HA tag that does not influence synaptic function (not shown).

**Table 1.** A 2D two-sample Kolmogorov-Smirnov test to identify differences between Frazzled LOF mutants and LOF flies driving different UAS Frazzled rescue constructs. A) The individual, independent comparisons made with the KS2D2S test revealed whether there are differences in the distributions of response latency averages and the proportion of volume taken up by gap junction antibody in GF terminals, with significant differences shown by asterisks (*). B) The KS2D2S comparing response frequencies to the proportion of volume taken up by gap junction antibody in GF terminals shows the same significant differences as the test in A, except for UAS-FraE1354A (see Results).

Two sample, 2D Kolmogorov-Smirnov test
|  | Sample 1<br>(n = 35 terminals) | Sample 2 | Sample 2<br>Terminals | P-value<br>(Response Latency vs GJ%) | P-value<br>(Response Frequency vs GJ%) |
| --- | --- | --- | --- | --- | --- |
| 1 | frazzled loss of function mutant<br>(w; fra<sup>3</sup>/fra<sup>4</sup>; R91H05::GFP/+) | frazzled heterozygous sibling<br>(w; fra<sup>3</sup>/CyO; R91H05::GFP/+) | n = 16 | 0.0001 | 0.0001 |
| 2 | frazzled loss of function mutant<br>(w; fra<sup>3</sup>/fra<sup>4</sup>; R91H05::GFP/+) | Full length Frazzled<br>(w; fra<sup>3</sup>/fra<sup>4</sup>; R91H05::GFP/UAS-Frazzled) | n = 38 | 0.0373 | 0.0076 |
| 3 | frazzled loss of function mutant<br>(w; fra<sup>3</sup>/fra<sup>4</sup>; R91H05::GFP/+) | Full length Frazzled<br>(w; fra<sup>3</sup>/fra<sup>4</sup>; R91H05::GFP/UAS-FraICD) | n = 23 | 0.0007 | 0.0004 |
| 4 | frazzled loss of function mutant<br>(w; fra<sup>3</sup>/fra<sup>4</sup>; R91H05::GFP/+) | Full length Frazzled<br>(w; fra<sup>3</sup>/fra<sup>4</sup>; R91H05::GFP/UAS-FraE1354A) | n = 13 | 0.0628 | 0.0404 |
| 5 | frazzled loss of function mutant<br>(w; fra<sup>3</sup>/fra<sup>4</sup>; R91H05::GFP/+) | Frazzled P1 domain deletion<br>(w; fra<sup>3</sup>/fra<sup>4</sup>; R91H05::GFP/UAS-FraΔP1) | n = 7 | 0.0013 | 0.0153 |
| 6 | frazzled loss of function mutant<br>(w; fra<sup>3</sup>/fra<sup>4</sup>; R91H05::GFP/+) | Frazzled P2 domain deletion<br>(w; fra<sup>3</sup>/fra<sup>4</sup>; R91H05::GFP/UAS-FraΔP2) | n = 20 | 0.0002 | 0.0054 |
| 7 | frazzled loss of function mutant<br>(w; fra<sup>3</sup>/fra<sup>4</sup>; R91H05::GFP/+) | Frazzled P3 domain deletion<br>(w; fra<sup>3</sup>/fra<sup>4</sup>; R91H05::GFP/UAS-FraΔP3) | n = 12 | 0.3085 | 0.5607 |
| 8 | frazzled loss of function mutant<br>(w; fra<sup>3</sup>/fra<sup>4</sup>; R91H05::GFP/+) | Wildtype unablated<br>(w; + ; R91H05::GFP/10x-UAS-GFP) | n = 8 | 0.0055 | 0.0098 |
| 9 | frazzled loss of function mutant<br>(w; fra<sup>3</sup>/fra<sup>4</sup>; R91H05::GFP/+) | Wildtype ablated<br>(w; + ; R91H05::GFP/10x-UAS-GFP) | n = 22 | 0.0282 | 0.0009 |
| 10 | frazzled loss of function mutant<br>(w; fra<sup>3</sup>/fra<sup>4</sup>; R91H05::GFP/+) | ShakB<sup>2</sup> heterozygous sibling ablation<br>(ShakB<sup>2</sup>/Fm6; + ; R91H05::GFP/R91H05::GFP) | n = 14 | 0.3688 | 0.0076 |
| 11 | frazzled loss of function mutant<br>(w; fra<sup>3</sup>/fra<sup>4</sup>; R91H05::GFP/+) | ShakB<sup>2</sup> ablation<br>(ShakB<sup>2</sup>/ShakB<sup>2</sup>; + ; R91H05::GFP/R91H05::GFP) | n = 10 | 0.0019 | 0.0003 |

Crosses can generate up to eight different genetic siblings. In each cross, two of these genetic siblings are the trans-heterozygous Frazzled LOF mutants. Heterozygous Frazzled LOF siblings also appeared as two genetic siblings and were phenotypically wild-type, and we used the w; *fra*^3^/Gla, Bc; R91H05::GFP-GAL4/Tb sibling as our central sibling control. This sibling is phenotypically wild-type and appears in all the crosses generated. Since the data collected from this sibling is consistent, we grouped the data collected across different crosses together for easier comparisons.

### Electrophysiology

Flies were embedded ventral side down in dental wax after anesthetizing with CO_2_. Physiological recordings were obtained bilaterally using glass microelectrodes placed directly into the tergotrochanteral jump muscle (TTM) and dorsal longitudinal muscle (Figure 2A, 2B; Allen et al., 2006; Allen & Godenschwege, 2010; Augustin et al., 2017; Orr et al., 2014). Each microelectrode was filled with O’Dowd’s Drosophila saline (Gu & O’Dowd, 2006). GFs were stimulated extracellularly using Grass stimulators with tungsten wire electrodes placed in the brain through the eyes, and a tungsten wire ground electrode was placed in the fly’s abdomen (Figure 2A, 2B; Allen & Godenschwege, 2010). An Axon Digidata 1440A Data Acquisition System was used to digitize the data, and recordings were collected using Clampex software (Molecular Devices). Muscle response latency was reported in milliseconds, and typical wild-type TTM latencies are <1.00ms. Response frequencies were obtained by initiating a train of 10 stimuli at 100 Hz every second for ten trains. These were reported as an average percentage of successful muscle responses per stimuli for each animal. Wild-type response frequencies are defined as ≥90% responses to train stimulation at 100Hz.

**Figure 2.**
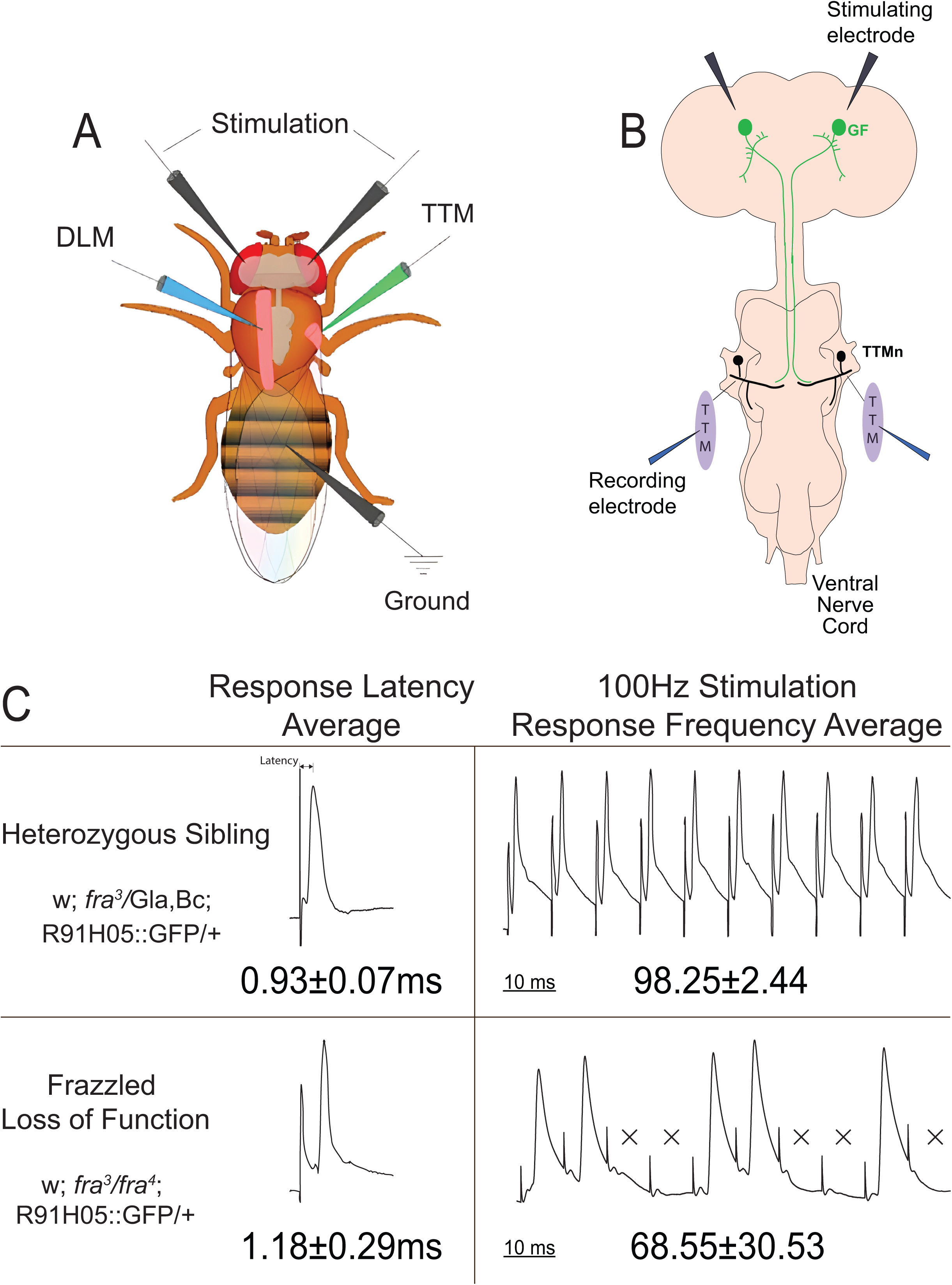
Recording from the Giant Fiber System. **A)** The Giant Fiber System (GFS) relays signals from the brain to the jump muscles in the thorax. We insert stimulating electrodes through the eyes to extracellularly stimulate the GF and insert recording electrodes at the jump muscle to record latency and response frequency for the circuit (see Methods) **B)** Schematic of the GF system, with recording electrodes placed extracellularly (TTM). **C)** Latencies are recorded ten times with one-second intervals between stimuli. Wild-type is defined by latency averages under 1.00ms. When we stimulate the GFs ten times in ten seconds at 100Hz, wild-type samples have consistent response frequency averaging over 90%. The upper panels are from heterozygous *frazzled* loss of function sibling samples and are wild-type, while the lower panels are *frazzled* loss of function mutants, which perform significantly worse in comparison.

Recorded responses to stimuli result from an individual GF terminal-TTMn connection. Therefore, each response is assigned to a terminal (left or right), and each terminal is treated as a separate unit for the purposes of statistics throughout the results, whether it belongs to a bilateral GF or a wild-type-appearing GF and the physiology and anatomy were assessed for each terminal unit separately.

### Immunohistochemistry

Flies were removed from dental wax after electrophysiology tests were concluded and dissected in phosphate-buffered saline (PBS). We opened the thorax and abdomen of the flies so that the ventral nervous system (VNS) was exposed (Boerner & Godenschwege, 2011), and the head case was also opened to reveal the brain. Samples were fixed in 4% paraformaldehyde for 45 minutes, washed in PBS overnight, then washed in a PBS 0.5% Triton X for 2 hours at room temperature. The GFs were labeled using primary antibodies: mouse anti-GFP (1:500; Invitrogen) and rabbit anti-shaking-B (1:500; Biomatik). Our anti-shaking-B was generated by Biomatik following a published peptide sequence (Phelan et al., 1996). Primary antibodies were prepared using PBS with Triton X 0.3% and bovine serum albumin (BSA) 3%. After incubating for two nights at 4°C, samples were washed for two hours in PBS, and secondary antibodies with conjugated fluorophores were applied overnight to detect proteins under confocal microscopy: goat anti-rabbit Cy2 (1:500) and goat anti-mouse Alexa Fluor 488 (1:500) at 4°C. Each sample was then washed in PBS for two hours and dehydrated in an ethanol series for ten minutes each in 50%, 70%, 90%, and 100% ethanol and mounted on glass slides filled with methylsalycylate and sealed with a cover slip for imaging.

### Imaging

Samples were imaged on a Nikon A1R laser scanning confocal microscope. Samples were magnified using a 60x oil immersion objective scanned at a pixel resolution of 1024 x 1024, and z-step sizes were 0.125µm. Maximum intensity projection images were created in Nikon NIS Elements software; contrast and brightness were adjusted to increase image quality.

To analyze the structure and content of the GFs, 3D volume renderings were generated in IMARIS software. Before rendering, background subtraction was applied to both channels. Rendering was performed with a 0.125 smoothing adjustment. Thresholding was applied separately on a sample-by-sample basis. Terminals were defined as the area of the GFs posterior to the peripherally synapsing interneuron (PSI) region (Figure 3 dotted lines). We applied a gap junction antibody mask to assess the volume in the GF terminals taken up by the gap junction antibody channel mask to show only the GF terminal content, excluding the antibody label outside the terminal.

**Figure 3.**
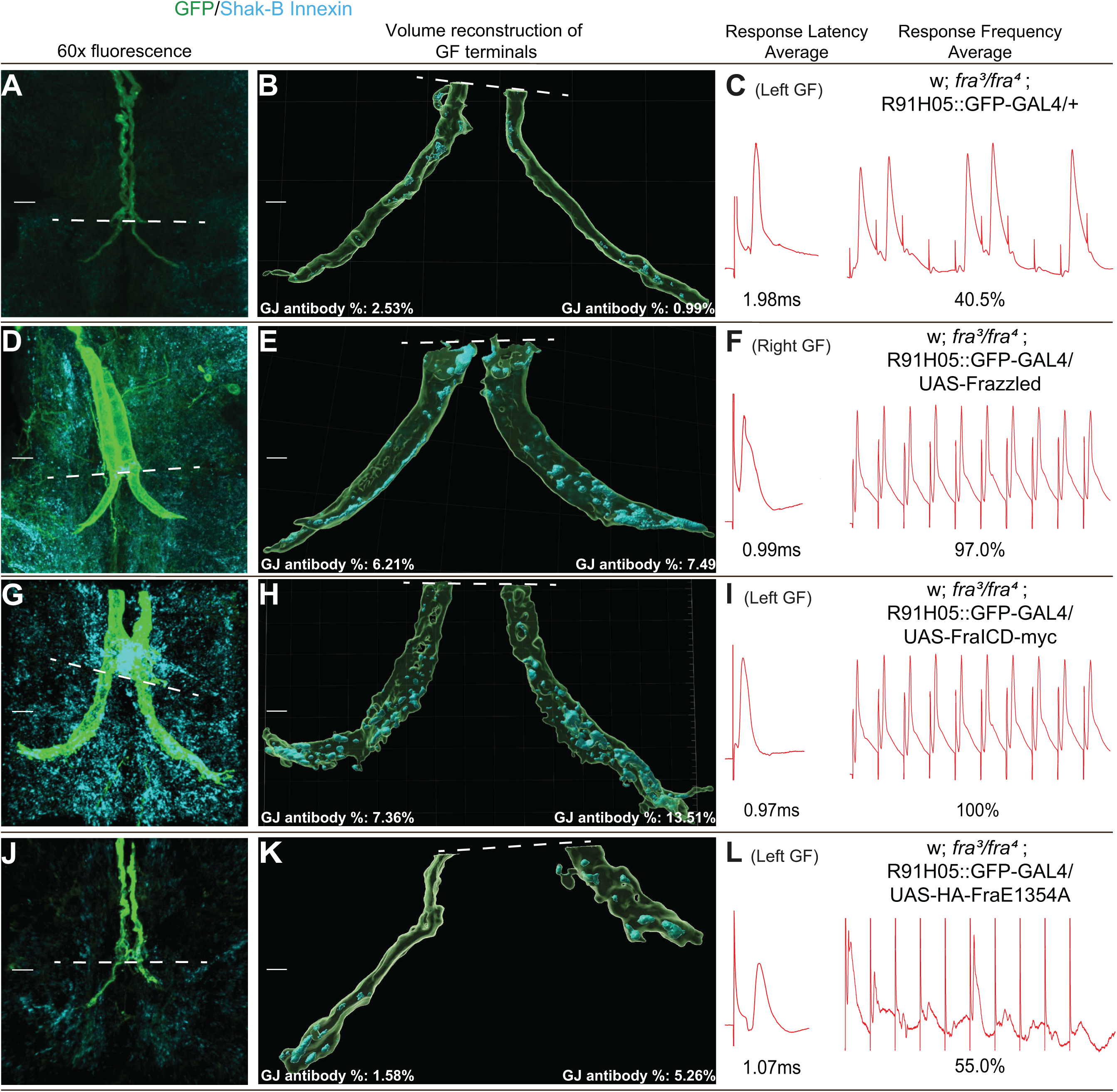
Structure of Giant Fiber axons when driving different UAS constructs in a Frazzled LOF background. Each row has images from an individual sample for a given genotype. First column (A, D, G, J): Compressed z-stack projection image of the Giant Fiber terminals labeled with GFP, and shaking-B(neural+16) gap junctions labeled with shaking-B antibody. The dotted line marks the PSI region and the anterior limit of the unit terminals. Second column (B, E, H, K): Volume reconstructions of the corresponding Giant Fiber terminals posterior to the PSI region. The reconstruction generates volumes for the GF terminals and the gap junction antibodies within the terminals, which are included for each sample. Third column (C, F, I, L): Genotype of the samples in each row and that sample’s latency and response frequency average.

### Statistical Analysis

A 2D two-sample Kolmogorov-Smirnov (KS2D2S) test was used to examine the differences between genotypes using latency and percentage of the synaptic terminal filled with antibody to the gap junction. The KS2D2S test was performed using Python’s NumPy and SciPy libraries and carried out using the scipy.stats library in Python, with a significance level (alpha) set at 0.05. Our hypothesis states that Frazzled has a transcriptional role in the GFs, and we show this by comparing response latencies, response frequencies, and the proportion of gap junction antibody volume to terminal volume from Frazzled LOF mutant flies to LOF flies driving different UAS-constructs. By comparing each experimental group to a control group using the KS2D2S test, we perform separate, independent tests and avoid raising the familywise error rate.

### Cell Ablation

In some *frazzled* mutants, only one GF grows from the brain to the thorax, and the GF usually splits to form a bilateral terminal. To investigate the bilateral terminal in a wild-type background, we ablated one GF and found that the remaining one formed a bilateral terminal (Boerner et al., 2024). The strength of the bilateral connections was assessed in the usual manner, and the percentage of the terminal occupied by antibodies to the gap junction was measured. After physiological recordings were performed, specimens were dissected to reveal the CNS, and GFs were dye-injected with a mix of TRITC-Dextran and Neurobiotin. The large Dextran molecule labels the GF axon and terminal but does not pass through gap junctions. Neurobiotin can cross the gap junctions between GF and TTMn, and a Neurobiotin signal was detected in the post-synaptic dendrite. This Neurobiotin dye-coupling confirmed intact gap junctions between the two neurons.

### Computational Modeling of the Giant Fiber

Our study utilizes a computational model composed of four interconnected neuronal compartments designed to investigate the effects of synaptic strength, gap junction conductance, and external stimuli on neuronal firing behaviors (Augustin et al., 2019; Follmann et al., 2015). The first compartment receives an excitatory stimulus, *I*_stim_, designed as a pulse current (10 Hz) combined with a zero-mean and unit-variance random Gaussian noise to mimic the variability observed in biological neuronal inputs. This external stimulation propagates through the other compartments (axial resistance) and can trigger action potentials in the postsynaptic neuron compartment, depending on the strength of the gap junction or chemical synapse that connects these neurons or the frequency and amplitude of the input signal.

### Neuronal Dynamics and Gating Variables

The model is based on Hodgkin-Huxley formalism, detailed through dynamic variables: membrane potential (*V*) and gating variables (*n*, *m*, and *h*). Ionic currents and synaptic inputs influence each compartment’s membrane potential (V) (*I_syn_*), defined by:

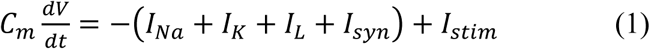

where *C_m_* denotes the membrane capacitance, *I_Na_*, *I_K_*, and *I_L_* represent the sodium, potassium, and leak currents. *I_syn_* and *I_stim_* symbolize synaptic and external stimulation currents. The gating variables (*x*=*n,m,h*) are defined by the following:

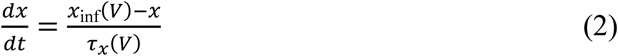

where the steady-state values of the gating variables for a given membrane potential *V* is given by:

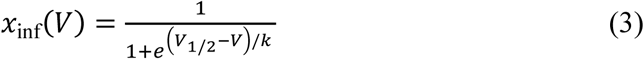

with *V*_1/2_ = −53 Mv and *k* = 15 for *n*, *V*_1/2_ = −40 Mv and *k* = 15 for *m*, and *V*_1/2_ = −62 Mv and *k* = −7 for h. The corresponding voltage-dependent time constants are governed by *τ_n_* = 1.1 + 4.7 exp(—), *τ_m_* = 0.04 + 0.46 exp(−—), and *τ_h_* = 1.2 + 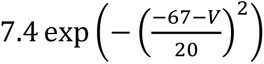. Re—l potentials are -85 mV for leak, -74 mV for potassium, and 65 Mv for sodium.

### Chemical and Electrical Signal Modelling

The model uses chemical (*I*_chem_) and electrical (*I*_gap_) synapses to simulate the rapid escape response of Drosophila Giant Fibers. The chemical synaptic current, *I*_chem,_ is defined as:

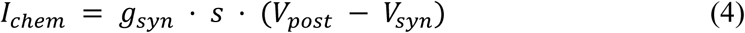

*g*_syn_ represents the maximum synaptic conductance, set to 0.1 in the model, *V*_post_ is the post-synaptic compartment’s membrane potential, and *V*_syn_ is the synaptic reversal potential, set to 0 mV, indicating an excitatory synapse. *s* represents the fraction of activated receptors or open synaptic channels at any given time. The dynamics of *s* is governed by the equation:

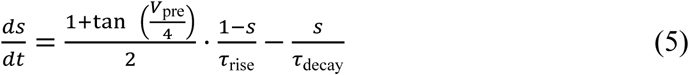

where *V*_pre_ is the membrane potential of the presynaptic compartment, *τ*_rise_ and *τ*_decay_ are the time constants for the synaptic activation and deactivation, respectively. In this model, *τ*_rise_ = 0.1ms and *τ*_decay_ = 3ms are used to simulate the synaptic response kinetics. For *I*_gap_, we assume *s*=1, and the electrical synaptic conductance value (*g_gap_*) is obtained from the percentage of terminal volume filled (*p*), allowing for a coupling strength that connects with the experimental samples through a sigmoid relationship:

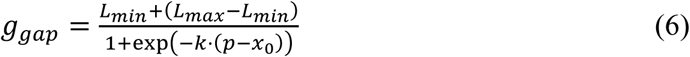

Here, *L_max_* and *L_min_* represent biologically grounded maximum and minimum gap junction conductance values, respectively. For values of *p*>10%, we assume *g*_gap_ = *L_max_*. The parameter *k* controls the steepness of the sigmoidal function, and *x*_0_ determines its midpoint. This formulation ensures that the conductance transitions smoothly from *L_min_* to *L_max_* as the percentage of terminal volume filled increases, capturing the biological variability in electrical coupling.

## RESULTS

### Axon pathfinding is disrupted in Frazzled LOF mutants

In adult *Drosophila*, GF axons grow posteriorly from the brain through the connective and along the midline to the ventral nerve cord to form an ipsilateral synaptic terminal with their TTMn partners in the second thoracic segment (Figure 1C). Previous studies in our lab found that GF axon pathfinding and synaptogenesis were disrupted in Netrin LOF and Frazzled LOF mutant flies (Orr et al., 2014). To examine the role of Frazzled in synaptogenesis more closely, we revisited this analysis and analyzed the GF system in Frazzled/DCC mutants and various UAS constructs (Figure 1I). In Figure 1A-G, we present the range of GF anatomical phenotypes seen in LOF flies (*fra*^3^/*fra*^4^) and quantify the distribution of phenotypes in Figure 1H. In approximately half of the LOF flies, the GFs appear wild-type, and each axon innervates the ipsilateral TTMn (n = 16 terminals; Figure 1A, 1H). In approximately the other half of the mutant flies, we find one phenotype never seen in wild-type animals, where one GF extends an axon within the brain, but the other never leaves the brain (n = 15 terminals; Figure 1D). The extended GF grew to the thorax and formed a bilateral terminal to innervate both TTMn targets (Figure 1B, 1F; Orr et al., 2014; Kennedy and Broadie., 2018; Boerner et al., 2024). The bilateral terminal is a direct result of one GF being lost in the brain and an indirect result of the Frazzled LOF mutation. Finally, the most severe pathfinding phenotype seen in a few LOF mutants occurred when both GFs grow long, twisted axons that never leave the brain, which we called “stuck-in-the-brain” (n = 4 GFs; Figure 1G). As a result, we record no motor responses to stimulation of these flies and can’t measure changes in gap junction antibody volume at the GF terminals. These flies are excluded from statistical analysis. The proportion of the various phenotypes for each genotype is recorded in Figure 1H.

### LOF mutants exhibit defective physiology

We next analyzed the physiology of the Frazzled LOF flies and compared them to sibling controls (w; *fra*^3^*/*Gla, Bc; R91H05::GFP-GAL4/Tb) that were used as the baseline control in all the subsequent crosses we generated. When we apply a supra-threshold stimulus to the GFs of Frazzled heterozygous siblings and record from the jump muscles, their response latencies average 0.93ms (SD = 0.07, n = 16 terminals; Figure 2C, Figure 4A). We stimulate the GFs 10 times in 100ms at 100Hz to determine the response frequency. Heterozygous siblings respond to each stimulus in 98.3% of the trials (SD = 2.44, n = 16 terminals; Figure 2C, Figure 4D). In contrast, Frazzled LOF mutant flies had poorer response latencies than their control siblings (1.18ms, SD = 0.30, n = 35 terminals; Figure 2C, Figure 4A) and reduced average response frequencies (68.55%, SD = 30.53, n = 35 terminals; Figure 2C, Figure 4D).

**Figure 4.**
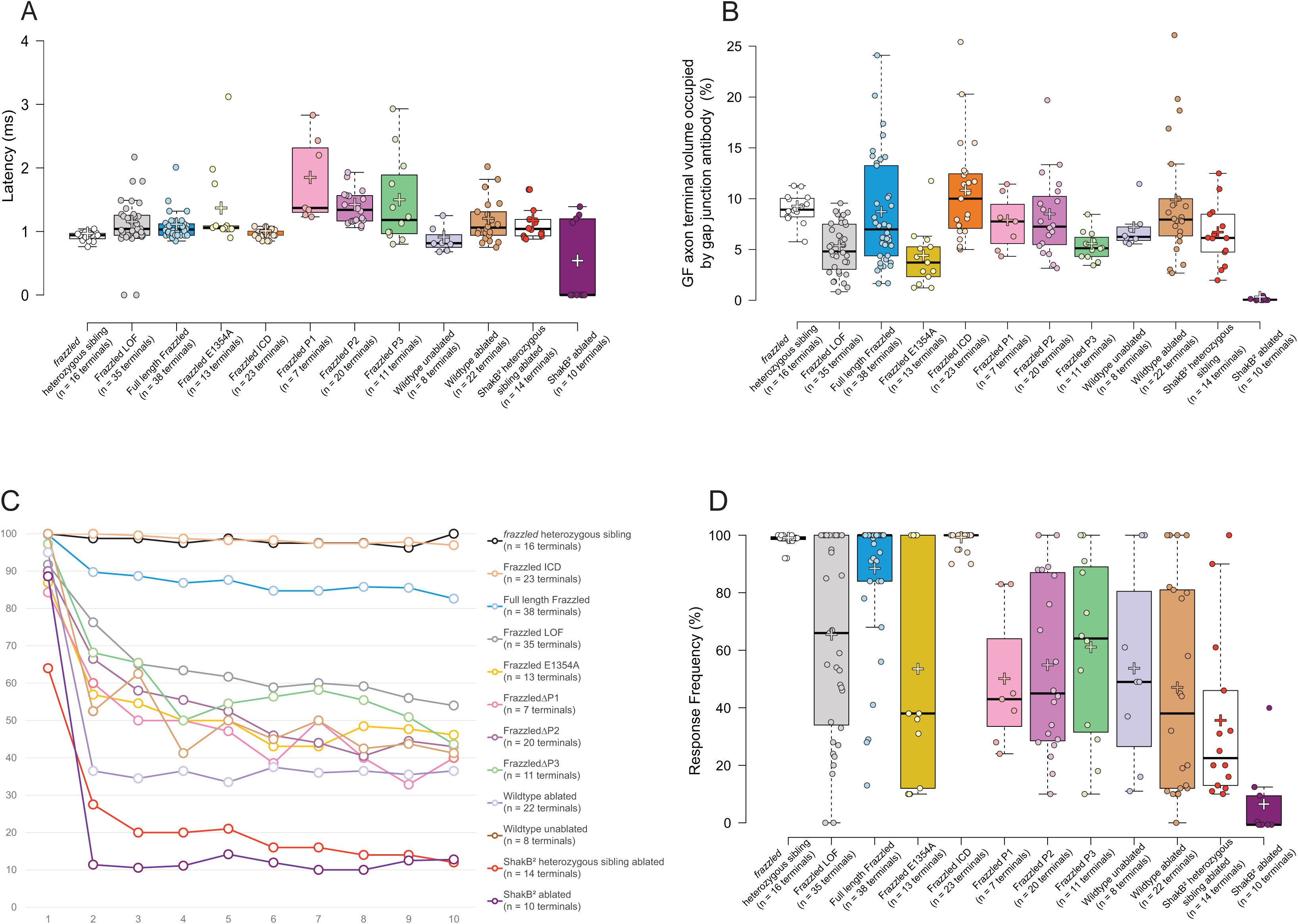
Comparison of latency, response frequency, and gap junction antibody proportion in GF terminals for all genotypes tested. **A)** Latency comparison for all genotypes tested. Each point is the latency average of ten trials from one side (unit terminal) of a specimen. **B)** We measured Giant Fiber terminal volume and gap junction antibody volume by generating a 3D rendering of Z stack confocal images using IMARIS software. We plotted the percent of a GF terminal occupied by gap junction antibody for each genotype tested. **C)** Left and right side response frequency average for each fly of each genotype tested. **D)** Response frequency comparison for all genotypes tested. Each point is the response average for a unit terminal stimulated 10 times at 100Hz. Center lines (A, B, D) show the medians; box limits indicate the 25th and 75th percentiles as determined by R software; whiskers extend 1.5 times the interquartile range from the 25th and 75th percentiles; crosses represent sample means; data points are plotted as open circles.

### Gap junctions are disrupted in Frazzled LOF mutants

Previous experiments quantified the gap junction immuno-histological label volume in the mutant and wild-type GFs (Orr et al., 2014). To revisit Orr’s results, we ordered a custom shaking-B innexin antibody and examined the distribution of gap junctions within the GFP-labeled terminals in mutant and control flies (see Figure 3 and Methods). We used generated 3D renderings of the confocal images and measured the volume of gap junction staining within the terminals. We defined the synaptic terminal as the region of the terminal posterior to the infra-medial bridge and the contact region for the GF and PSI (peripheral synapsing interneuron; Figure 3B dashed white lines). The differences are visible in the volume renderings, such as Figures 3B and 3E. In control siblings, gap junction antibody occupies 9.04% (SD = 1.33, n = 16 terminals; Figure 4B) of GF terminal volume on average, and in the Frazzled LOF mutants, gap junction antibody occupies 5.31% on average (SD = 2.40, n = 35 terminals; Figure 4B). Overall, the results are similar to those of Orr (2014), who used slightly different methods and reported that 10.93% of the control terminal volume was occupied by gap junctions and 3.00% of the terminal in Frazzled LOF specimens.

Since latency and gap junction antibody volume have been linked directly in the GFS (Augustin et al., 2017), we compared the latency averages and the proportion of volume taken up by gap junction antibody in GF terminals for each sample. To compare the various genotypes, we ran a two-dimensional, two-sample Kolmogorov-Smirnov (KS2D2S) test comparing the latency (and separately, the response frequency) averages to the average proportion of gap junction antibody volume in unit terminals. In Table 1 we show the results of the KS2D2S test done comparing the Frazzled LOF mutant (Sample 1 column, *fra*^3^/*fra*^4^) and all our tested UAS constructs (Sample 2 column). We listed the number of terminals compared for each UAS construct in a separate column. For example, in the first row of Table 1 we compared Frazzled LOF mutants and their heterozygous siblings (*fra*^3^/+). We found significant differences in the distributions of latency averages to the proportion of volume taken up by gap junction antibody in GF terminals for our samples (P = 0.0001, Table 1). Similarly, response frequency and gap junction volume were significantly different (P = 0.0001, Table 1). These results strongly suggest that the Frazzled LOF is associated with gap junction loss at the GF terminals.

### Bilateral terminals are an indirect phenotype of Frazzled LOF

Bilateral terminals appeared in most genotypes in these experiments (Figure 1B, 1H). This phenotype was especially interesting because it implies a competition between GFs for synaptic space on the postsynaptic motor neurons. It suggests that Fra/DCC might regulate the competition for synaptic space on the postsynaptic neurons. However, a detailed analysis of the phenomena demonstrated that the phenotype indirectly affects the pathfinding error, and the same phenotype can be induced by the ablation of one GF in wild-type animals (Boerner et al., 2024).

To better understand the bilateral terminals in the context of Frazzled, we created bilateral terminals in wild-type animals by laser ablation of one GF and analyzed the remaining GF, which nearly always (13/15) produced a bilateral terminal (Boerner et al., 2024). We collected the same data for these bilateral terminals: latency, response frequency, and proportion of the synaptic terminal occupied by the antibody to gap junctions and used the KS2D2S test to compare the *frazzled* mutant with bilateral terminals to the wild-type specimen (Table 1). Since the recordings were obtained from individual motor neurons innervated by half of the bilateral terminal, we treated each half terminal as a unit. We measured the latency, response frequency, and gap junction antibody volume. Ablated bilateral terminals created in a wild-type background exhibit latency (1.16ms, SD = 0.33, n = 22 terminals; Figure 4A) and response frequency averages (46.23%, SD = 36.15, n = 22 terminals; Figure 4D), but wild-type proportions of gap junction antibody volume averages in the terminals (9.65%; SD = 5.76, n = 22 terminals; Figure 4B).

When we separate the terminals found in Frazzled LOF specimens into two groups (wild-type-appearing and bilateral), they are functionally indistinguishable. The wild-type-appearing unit terminals exhibit a latency of 1.14ms (SD = 0.32, n = 20 terminals; Figure 4A) and response frequency of 79.60% (SD = 25.73, n = 20 terminals; Figure 4D), while the unit terminals from bilateral GFs exhibit a latency of 1.23ms (SD = 0.25, n = 13 terminals; Figure 4A) and a response frequency of 51.54% (SD = 29.50, n = 13 terminals; Figure 4D).

In a Frazzled LOF wild-type-appearing unit terminal and bilateral unit terminal, both have similar proportions of gap junction antibody volume to terminal volume averages (7.30µm^3^, SD = 4.80, n = 20 terminals; 8.24µm^3^, SD = 8.34, n = 16 terminals, data not shown). This suggests that bilateral GFs produce double the amount of gap junction compared to wild-type-appearing GFs. Similarly, wild-type-appearing and bilateral terminals exhibit fewer gap junctions in Frazzled LOF terminals than control ones (Figure 4B). On average, gap junction antibody occupied 5.54% of wild-type-appearing GF terminals (SD = 2.29, n = 20 terminals) and 4.61% of the bilateral unit terminals (SD = 2.51, n = 16 terminals). Since the unit terminals from bilateral GFs are so similar to wild-type-appearing GF terminals, we kept the unit terminals from both groups together when we analyzed the results for all the genotypes throughout the remainder of the results.

### Full-length UAS-Frazzled rescues the Giant Fiber electrical synapse

To refine our idea that Frazzled controls the assembly of the electrical synapse, we dissected the Frazzled receptor using the UAS-GAL4 system. We drove several different UAS-Frazzled constructs (Figure 1I; Neuhaus-Follini & Bashaw, 2015) in a Frazzled LOF background and assayed the ability of a given construct to rescue the LOF phenotypes.

First, we drove full-length UAS-Frazzled in the GFs of Frazzled LOF mutants using the GF driver R91H05-GAL4 recombined with green fluorescent protein (GFP) to label the GF axons (w; fra3/fra4; R91H05::GFP-GAL4/UAS-Frazzled, Figure 1I; Borgen et al., 2017; Jenett et al., 2012; Orr et al., 2014). We analyzed the physiology of 20 Frazzled LOF flies where UAS-Frazzled was driven in the GFs and found they had response latencies averaging 1.07ms (SD 0.20, n = 38 terminals; Figure 4A) and response frequencies averaging 87.17% (SD 22.90, n = 38 terminals; Figure 4D). We generated 3D volume renderings to assess the volume of anti-shaking-B gap junction antibody present within each unit terminal of LOF flies driving UAS-Frazzled in the GFs (Figure 3D, 3E). We compared response latency and the proportion of volume taken up by gap junction antibody in GF terminals of UAS-Frazzled and Frazzled LOF mutants. We found a significant difference between the distributions for our samples using a KS2D2S test (P = 0.0373, Table 1). A similar difference was found when comparing response frequency and proportion of volume taken up by gap junction antibody in GF terminals (P = 0.0076, Table 1). This rescue of synaptic structure and function strongly supports our hypothesis that UAS-Frazzled regulates synaptogenesis in the GFs.

### The ICD of Frazzled rescues GF synaptic structure and function

Frazzled/DCC’s intracellular domain (ICD) is known to be involved in several intracellular signaling pathways (Boyer & Gupton, 2018), so we next attempted to rescue the LOF phenotypes with the Frazzled-ICD. We analyzed the GF anatomy of 23 flies and found a partial rescue of the axon guidance phenotypes. Seventeen of the flies had wild-type appearing GFs, six had one GF stuck in the brain, and none had both GFs stuck in the brain (Figure 1H). We analyzed the latency, response frequency, and proportion of volume taken up by gap junction antibody in GF terminals of 12 flies and found they exhibited wild-type responses and had an average response latency of 0.97ms (SD 0.06, n = 23 terminals; Figure 4A) and a response frequency average of 98.40% (SD 3.16, n = 23 terminals; Figure 4D). A KS2D2S test between the Frazzled LOF mutants and mutants driving Frazzled ICD found a significant difference between the distributions of response latency and response frequency averages to the proportion of volume taken up by gap junction antibody in GF terminals (P = 0.0007, P = 0.0004, Table 1). These findings demonstrate that the Frazzled ICD nearly completely rescues all the LOF mutant phenotypes independent of Frazzled’s extracellular domain. We suspect the difference between UAS-Frazzled and UAS-FraICD comes from a rate-limiting step in Frazzled processing, where UAS-Frazzled undergoes additional molecular processes leading to extracellular cleavage to release its intracellular domain. UAS-FraICD generates intracellular domains that may skip these processes, leading to stronger rescue effects.

### Deleting the conserved Intracellular Domains (P1, P2, and P3) alters rescue outcomes

To further investigate what part of the intracellular domain is essential for GF structure and function, we tested three constructs that each express full-length Frazzled, each missing one of its highly conserved intracellular domains: UAS-FraΔP1, UAS-FraΔP2, and UAS-FraΔP3 (Figure 1I; Neuhaus-Follini & Bashaw, 2015). Driving these constructs individually in Frazzled LOF backgrounds revealed the rescue of distinct GF terminal phenotypes for the different domains. For UAS-FraΔP1, we analyzed 13 flies and found none had wild-type appearing terminals, six had bilateral terminals, and seven had both GFs stuck in the brain (Figure 1H). Latencies averaged 1.80ms (SD = 0.61, n = 7 terminals; Figure 4A), response frequencies averaged 54.05% (SD = 29.14, n = 7 terminals; Figure 4D), and gap junction antibody made up 7.65% of the unit terminal volume on average (SD = 2.53, n = 7 terminals; Figure 4B). Here, the KS2D2S showed that the results for latency, response frequency, and gap junction volume were different from LOF, implying a partial rescue of the gap junctions (P = 0.0013, P = 0.0153, Table 1) Similarly, For UAS-FraΔP2, we analyzed 10 flies and found two had wild-type appearing terminals, eight had bilateral terminals and none had both GFs stuck in the brain (Figure 1H). Latencies averaged 1.40ms (SD = 0.25, n = 20 terminals; Figure 4A), response frequencies averaged 54.05% (SD = 29.14, n = 20 terminals; Figure 4D), and gap junction antibody in the unit terminals occupied 8.26% of the unit terminal volume on average. (SD = 3.85, n = 20 terminals; Figure 4B). Similar to ΔP1’s KS2D2S result, flies driving UAS-FraΔP2 also showed significant differences compared to LOF flies (P = 0.0002, P = 0.0054, Table 1). These results suggest a partial rescue of the gap junctions using these constructs (UAS-FraΔP1 and UAS-FraΔP2).

In contrast to UAS-FraΔP1 and UAS-FraΔP2, we analyzed 20 UAS-FraΔP3 flies and found seven had wild-type appearing terminals, eight had bilateral terminals, and five had both GFs stuck in the brain (Figure 1H). Here, response latencies averaged 1.46ms (SD = 0.69, n = 11 terminals; Figure 4A), response frequencies averaged 60.00% (SD = 31.67, n = 11 terminals; Figure 4D) and gap junction antibody occupied 5.37% of the unit terminal volume on average (SD = 1.41, n = 11 terminals; Figure 4B). The KS2D2S test showed UAS-FraΔP3 is not different from LOF, demonstrating the absence of any rescue effect on the synapse (P = 0.3085, P = 0.5607, Table 1). Taken together, the data suggest that the P3 domain of Frazzled is required for regulating gap junctions in the GF while P1 and P2 are not necessary.

### The Frazzled transcriptional activation domain is required to rescue synaptic function

To further dissect Frazzled’s ICD, we drove a full-length Frazzled construct containing a point mutation in the P3 domain in a LOF background (Figure 1I). This ICD mutation (FraE1354A) is known to prevent transcription of commissureless, a midline guidance protein expressed during larval development (Neuhaus-Follini & Bashaw, 2015). When we analyzed the GF terminals of LOF flies driving UAS-HA-FraE1354A, we found that this construct rescued neither axon guidance nor synaptogenesis. Three flies exhibit wild-type appearing terminals, six flies with bilateral terminals, and five with both GFs stuck in the brain (Figure 1H). When we analyzed the physiology of UAS-HA-FraE1354A flies, we found they had an average latency of 1.33ms (SD 0.60, n = 13 terminals; Figure 4A), a response frequency average of 52.69% (SD 38.72, n = 13 terminals; Figure 4D) and gap junction antibody occupied 4.18% of the unit terminal volume on average (SD = 2.75, n = 13 terminals; Figure 4B).

When we compared the distributions of response latency and response frequency averages each to the proportion of volume taken up by gap junction antibody in unit terminals of UAS-HA-FraE1354A flies and Frazzled LOF mutants with a KS2D2S test, we found no significant difference between the response latency and average proportion of gap junction antibody volume to terminal volume but did find a difference between response frequency and proportional gap junction antibody (P = 0.0628, P = 0.0404, Table 1). The average latency, response frequency and proportion of gap junction antibody in unit terminals are worse when we drive UAS-HA-FraE1354A flies in a Frazzled LOF background compared to Frazzled LOF alone. Taken together, this finding suggests that UAS-HA-FraE1354A produces a dominant negative effect on the Frazzled LOF mutant, strongly supporting our hypothesis that Frazzled’s transcriptional activation domain is required to rescue gap junctions in a Frazzled LOF mutant background.

### Modeling the role of gap junctions in synaptic function

As the amount of gap junction was reduced in the experimental animals, the latency increased and response frequency decreased (Figure 4A, 4C). Since gap junctions are not typically use-dependent, the increased rate of response decrement was an enigma. To explore the role of gap junctions in response decrement, we created a model of the GF-TTMn circuit and simulated the amount of gap junctions available at the virtual synapse in a manner that parallels the experimental data with *frazzled* mutants. A similar circuit, which we adapted, was previously modeled in a slightly different context to look at gap junctions during aging (Augustin et al., 2019). Our primary goal was to determine how gap junction-controlled circuits function, namely the latency and response frequency of the GF-TTMn synapse in our various genotypes. The most prominent feature of the function is the dramatic response decrement induced in multiple genotypes exhibiting different levels of gap junction in the terminals (Figure 4C).

We first developed a compartment model of the GFs that mimicked the signal-propagating properties of the axon and then modeled the electrochemical synapse to connect the model GF to a postsynaptic compartment. We used parameters found in the Augustin (2019) model to represent the features of our compartment model GF. We then reduced the amount of gap junction in the model in a manner that paralleled the results from our various *frazzled* genotypes. The basic properties of the model are shown in Figure 5. A signal is introduced in the furthest compartment from the synapse (blue) and propagates into connected compartments until it reaches the synapse (green), where the electrochemical dynamics determine if an action potential is produced.

**Figure 5.**
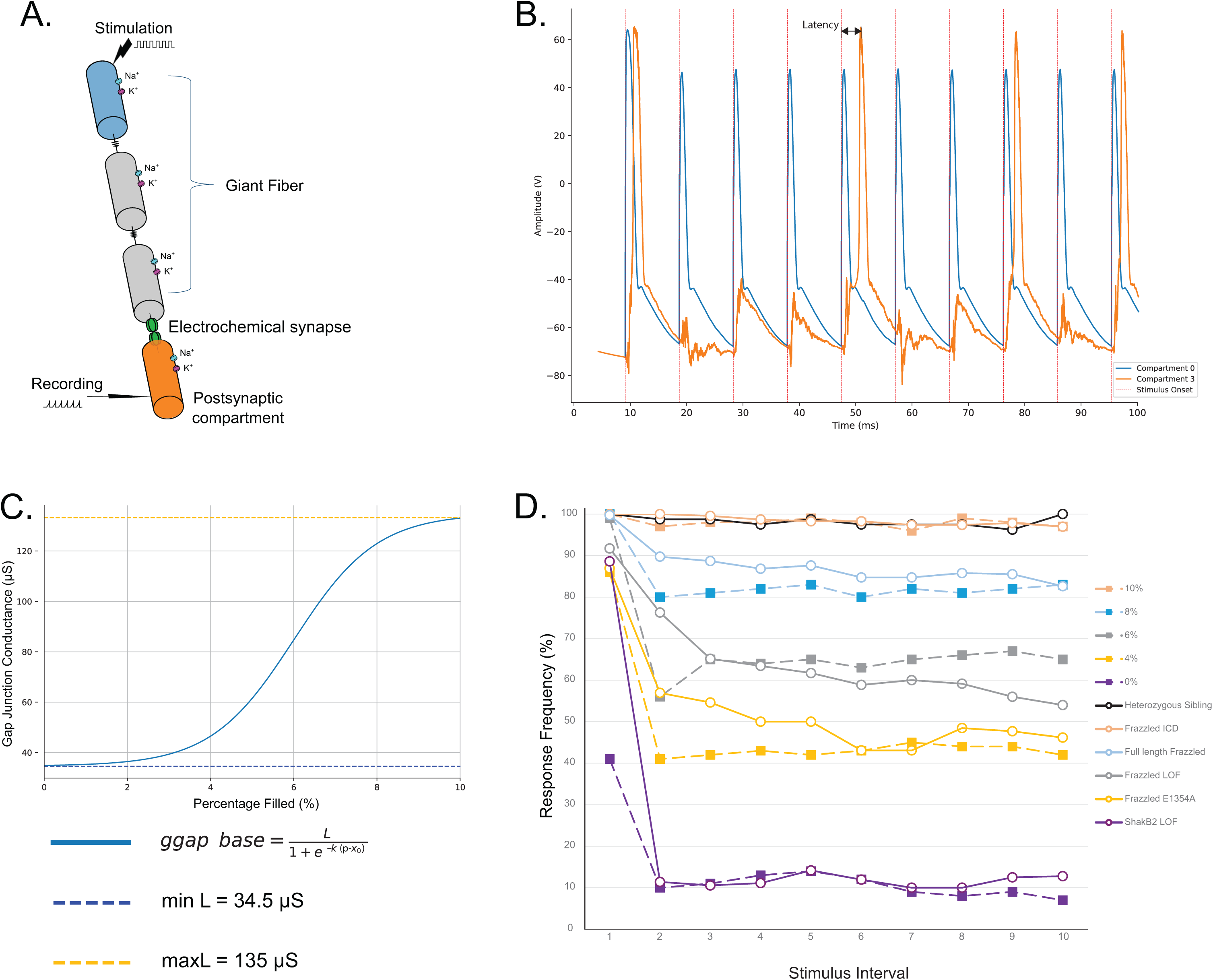
Modeling the GF-TTMn synapse effectively simulates responses to stimulation when altering gap junction proportions in terminals. A) Our model uses compartments to simulate the Giant Fiber system. A stimulus is introduced in compartment 0 and is propagated through subsequent compartments that make up the GF model. Between compartments 2 and 3, we placed a simulated electrochemical synapse and varied the amount of gap junction present at the synapse to alter responses to stimulation. An action potential is then produced in compartment 3, where response latency and frequency are recorded. **B)** Our model can mirror experimental responses accurately. The recordings from compartment 3 can be seen in orange. Latency is calculated as the difference in time between the peaks of compartment 0, shown in grey, and compartment 3. Response frequency is measured by the number of action potentials recorded after 100Hz stimulation. **C)** Gap junction conductance in the model is modulated by a sigmoid transformation. In wild-type flies, the amount of gap junction protein present in the GF terminals is on average 10% of the total volume of the terminal. In a *shak-B^2^* mutant, gap junctions are absent from the GF-TTMn synapse, and the GF relies solely on the chemical synapse for signaling (Boerner et al., 2024). Gap junctions are naturally lost as the fly ages, so we set our limits of gap junction conductance following parameters found in Augustin (2019) that follow age-related decreases in conductance and added an exception for zero gap junctions to lead to zero conductance. We used a sigmoid equation as a simplification of the Boltzmann equation since we want to look at the effects of varying gap junction proportions on responses and not channel dynamics. **D)** We find that our model (dashed lines) can closely mimic the response decrement profile of our *frazzled* mutant flies (solid lines).

The amount of gap junction observed in our experimental data was used to set the range of gap junction function in the model. The volume of anti-gap junction antibody in a control terminal is approximately 10% of the terminal volume and 5.31% in Frazzled LOF (Figure 4B). We used *shak-B^2^* flies as the lower boundary of gap junction antibody, which was shown to be < 1% of the terminal ((Boerner et al., 2024). We applied a sigmoid transformation, as opposed to a Poisson-Boltzmann transformation, to simplify the conductance-per-gap junction amount calculation and set the upper and lower limits of conductivity based on parameters found in the Augustin (2019) model.

The model showed latency increases as the amount of gap junction was reduced (Figure 5B). This was expected and consistent with the Augustin (2019) data. What was somewhat surprising was that the response frequency decreased as the proportion of gap junction was reduced (Figure 5D, dashed lines). Since gap junctions are not typically thought to show activity dependence, the source of this response decrement was unclear and not explored by Augustin (2019). Using the volume of gap junction antibody as a proxy for gap junction function, we simulated the model’s response at different proportions of gap junction seen in various genotypes. The model accurately simulates the response decrement as shown by the data from multiple genotypes (Figure 5D). Reducing the gap junction proportion to zero dramatically reduced the response latency and frequency, leaving a very weak response, as seen in mutants with no gap junctions (*shak-B^2^*). Removal of the chemical synapse in the model when the gap junctions were between 0 and 10% did not significantly alter synaptic function, as in the wild-type animal.

## DISCUSSION

The results confirm our hypothesis that Frazzled regulates synaptogenesis in the Giant Fibers by regulating gap junctions. Specifically, we show that Frazzled/DCC regulates the presynaptic gap junctions shaking-B(neural+16) that make up the electrical synapse of the GF. We quantified synaptogenesis by using response latency, response decrement, and the amount of gap junction antibody in the GF terminals. By combining electrophysiological data and confocal imaging, we are the first to show how Frazzled regulates electrical synapses through transcription. Simulation studies demonstrated how this control of gap junctions regulated the latency and response decrement of the synapse.

### Frazzled regulates the synaptic structure and function in Giant Fibers

In *frazzled* LOF mutant flies, the GFs exhibit long response latencies and rapid response decrement due to the reduced density of gap junctions (Figure 4A, 4C, Table 1, Orr et al., 2014). We focused on the presynaptic side of the GF-TTMn electrical synapse, which is assembled from the shaking-B(neural+16) isoform (Phelan et al., 2008). Our findings show that antibodies to gap junctions make up 9.04% of GF terminal volume in control siblings and 5.14% of GF terminal volume in LOF mutants (Figure 4B).

### The Frazzled ICD rescues synaptogenesis

To analyze Frazzled’s synaptogenic role in more detail, we attempted to rescue synaptic function in LOF mutants by driving various fragments of the Frazzled receptor in the LOF background (Figure 1I; Neuhaus-Follini & Bashaw, 2015). The most dramatic result occurred when we drove a UAS construct of the highly conserved Frazzled intracellular domain (UAS-FraICD, Figure 1I). This construct rescued most of the LOF phenotypes, including all the synaptogenic phenotypes (Figure 1H, Figure 4, Table 1). The results show response latency averages, response decrement, and the proportion of volume occupied by gap junction antibody in GF terminals flies carrying the rescue construct are restored to wild-type levels (Figure 4)

The dramatic rescue effects of the intracellular domain led us to examine other variations of the Frazzled receptor focused on the ICD to better understand its function in synapse formation. The intracellular domain contains a transcriptional activation domain overlapping the Frazzled P3 domain (Neuhaus-Follini & Bashaw, 2015). To examine the role of the P3 domain in a synaptic context, we drove a full-length Frazzled construct, lacking the P3 domain (UAS-FraΔP3, Figure 1I) in the Frazzled LOF background and assessed synaptic structure and function. This construct did not rescue the synaptic defects in the LOF mutants (Table 1).

To extend this idea, we drove a full-length Frazzled construct with a point mutation in the highly conserved P3 domain that is known to silence the transcriptional activation domain in the context of axon pathfinding in embryos (UAS-FraE1354A, Figure 1I; Neuhaus-Follini & Bashaw., 2015). In this rescue experiment, the physiology and proportion of gap junction antibody in terminals were not different from those seen in Frazzled LOF mutant flies (Table 1). These two constructs are known to affect transcription in embryos (UAS-FraΔP3 and UAS-FraE1354A; Neuhaus-Follini & Bashaw, 2015), and we suggest that Frazzled’s transcriptional activation domain is required for gap junction formation in the GFs. We hypothesize that Frazzled’s transcriptional activation domain regulates transcription of the innexin shaking-B(neural+16).

### The role of Frazzled/DCC in GF Axon Guidance

Our primary interest in this work was the control of synapse formation and function. However, Frazzled/DCC is a well-known axon guidance molecule, and we observed several disruptions of axon guidance linked to Frazzled/DCC. In the *fra*^3^/*fra*^4^ LOF animals, nearly half of the GFs exhibit severe guidance defects, grow randomly in the brain, never reach the thorax, and do not make synaptic connections with their normal targets. When full-length Frazzled was driven in the LOF mutants, the trends in the rate of GF axon pathfinding errors decreased overall but were not significantly different from Frazzled LOF mutants. We especially noted no change in the frequency of stuck-in-the-brain pathfinding errors. In contrast to the full-length construct, driving the ICD in the LOF mutants nearly completely rescued these stuck-in-the-brain defects. The strong effects of the ICD led us to examine the role of the ICD in axon guidance in more detail.

### The role of the P1, P2, and P3 domains in axon guidance

After testing the full-length Frazzled construct (UAS-Frazzled, Figure 1I) in a LOF background, we sought to dissect the ICD to understand how the highly conserved domains within the ICD function in axon pathfinding. We drove full-length constructs lacking one of the three domains (ΔP1, ΔP2, or ΔP3) in Frazzled LOF mutants (Figure 1I). The constructs lacking the ΔP1 and ΔP2 domains did not rescue axon guidance defects but did rescue the physiology and gap junction antibody proportion. In contrast, the ΔP3 construct did not rescue the axon guidance phenotypes nor the synaptic defects (Figure 1H, Table 1).

### A computational model of the GF-TTMn synapse

We developed our computational model of the GFS in Drosophila as a tool to assess synaptic function (Figure 5A, 5B). We used a conductance-based, multicompartmental model with ionic currents derived from the GF (Augustin et al., 2019). Despite our morphological and ionic modifications to the model, we validated the observed responses across a series of variations available in Augustin (2019) and obtained close agreement.

The model allows us to generate a proof of concept for our study of the specific role of gap junctions while keeping other features fixed, offering a robust framework for understanding the mixed electrochemical synapse. One of the primary strengths of the model is its striking similarity to experimental data. A novel aspect of our model is the sigmoidal function that dynamically adjusts gap junction conductance based on the volume of the terminal occupied by gap junctions (Figure 5C). This approach mirrors biological variability and allows for precise control of synaptic strength, making the model particularly suited for studying the effects of reduced gap junctions, as observed in *frazzled* LOF mutants. The rationale behind sigmoid functions comes from the previous use of Boltzmann functions to model the typical non-linear conductance change between two values (Moreno et al., 1995; Valiunas et al., 1999). When we run the model using gap junction proportion values close to those found in our experimental genotypes, translated by a sigmoid function (Figure 5C), we find that the model’s response frequency closely mirrors our data (Figure 5D).

The model highlights the strong relationship between response frequency and gap junction proportions in the GF terminal. We ran the model from 0-10% gap junction antibody proportion in the GF terminal compartment and generated a range of response frequency curves from the data. We find that these curves can closely predict the gap junction antibody proportion of flies given their response frequency data. To test this, we looked at response frequency data from flies expressing *shak-B^2^*, a mutation that silences gap junctions in the GFS (Phelan et al., 2008). In this experiment, flies were partially rescued by driving shaking-B (neural+16) protein in the *shak-B^2^* background. By comparing the response frequency data from these experiments to our model’s range of response frequency curves, we predict that *shak-B^2^* mutant flies using A307 to drive shaking-B(neural+16) have 6.6% of their terminal volume occupied by gap junctions, while mutant flies using C17 to drive shaking-B(neural+16) have 5.4% (Phelan et al., 2008).

The model incorporates periodic inputs with Gaussian noise to mimic biological variability, providing a realistic simulation of external stimuli affecting the GFs and synaptic failure in its transmission. However, the model also has limitations that warrant consideration. One such limitation is the simplification of certain aspects of synaptic dynamics, such as terminal morphology. The model is designed under the implication that a wild-type GF terminal connection to the TTMn is made. When terminals are bilateral or otherwise deformed, the distribution of gap junctions and other synaptic proteins changes. We can model such changes indirectly changing signaling dynamics in the model, but the exact changes happening *in vivo* are not fully understood. However, the flexibility of the model allows for sub-compartments to be created, mimicking bilateral morphology. Each sub-compartment can then be modified with their own gap junction proportions and chemical signaling changes to reflect the bilateral nature of the GF terminal. We may still learn something from these future expansions of the model to include bilateral terminals. Future refinements to the model will allow us to simulate synaptic connections beyond the traditional GFS and will let us build a model of the escape reflex from sensory neuron to motor neuron and every interneuron in between.

## Acknowledgments

This work was supported by grants from the Jupiter Life Science Initiative. We thank Dr. Greg Bashaw for his fly lines used in the work.

